# Uncertainty in contextual and kinematic cues jointly modulate motor resonance in primary motor cortex

**DOI:** 10.1101/293977

**Authors:** Andreea Loredana Cretu, Kathy Ruddy, Maria Germann, Nicole Wenderoth

## Abstract

Contextual information accompanying others’ actions modulates “motor resonance”, i.e. neural activity within motor areas that is elicited by movement observation. One possibility is that we weight and combine such information in a Bayesian manner according to their relative uncertainty. Therefore, contextual information becomes particularly useful when others’ actions are ambiguous. It is unclear, however, whether this uncertainty modulates the neural activity in primary motor cortex (M1) during movement observation. Here we applied single-pulse transcranial magnetic stimulation (TMS) while subjects watched different grasping actions. We operationalized motor resonance as grip specific modulation of corticomotor excitability measured in the index (FDI) versus the little finger abductor (ADM). We experimentally modulated either the availability of kinematic information (Exp. 1) or the reliability of contextual cues (Exp. 2). Our results indicate that even in the absence of movement kinematics, reliable contextual information is enough to trigger significant muscle-specific corticomotor excitability changes in M1 (p<.0001) which are strongest when both kinematics and contextual information are available (p<.005). These findings suggest that bottom-up mechanisms that activate motor representations as a function of the observed kinematics, and top-down mechanisms which activate motor representations associated with arbitrary cues converge in M1 in a statistically optimal manner.

## 1. INTRODUCTION

Neurophysiological studies have consistently demonstrated that observing movements performed by another individual triggers motor resonance, (i.e. activity within the mirror neuron system (MNS)) which responds similarly to action execution and action observation. MNS activity is believed to converge to cortical motor areas where it modulates activity in primary motor cortex (M1) such that changes in corticomotor excitability provide a readout of the neural net effect caused by motor resonance (for a review, see Naish et al., 2014). While it is well established that motor resonance is evoked by the observed kinematics, recent evidence indicates that higher-order information, such as prior knowledge or contextual cues, can facilitate intentional inference, even in the case of noisy or incomplete kinematics (e.g., Amoruso et al. 2016; Hudson et al. 2016). This is in line with predictive coding principles which state the importance of combining prior knowledge with sensory inputs for action understanding (Kilner, Friston, & Frith, 2007).

It has been suggested that contextual information can directly modulate activity in motor cortex, a finding that has mainly been derived from paradigms testing (i) anticipatory neural activity in motor cortex which precedes kinematic movement stimuli (e.g., Kilner et al. 2004a; Borroni et al. 2005); or (ii) neural activity when the observed kinematics and the contextual information are either congruent or incongruent (e.g., Janssen et al. 2015; Riach et al. 2018). More specifically, predictable contexts have been shown to increase the readiness potential in motor cortex (an index of motor preparation, Kilner et al. 2004a; Fontana et al. 2012) and increase corticomotor excitability specific to the muscles involved in the predicted actions (Alaerts, de Beukelaar, Swinnen, & Wenderoth, 2012; de Beukelaar, Alaerts, Swinnen, & Wenderoth, 2016; Sartori, Betti, Chinellato, & Castiello, 2015). These findings suggest that mirroring responses can be measured even prior to observing the kinematics of someone else’s movements. However, once the action unfolds, this pattern is updated to match the specific properties of observed movement kinematics (Cavallo, Bucchioni, Castiello, & Becchio, 2013). Particularly, if there is a mismatch between the predicted and observed movements, motor resonance will correspond to the muscles involved in performing the action, however, the effect will be lower compared to when no mismatch exists (Alaerts, Swinnen, & Wenderoth, 2010; Gueugneau, Mc Cabe, Villalta, Grafton, & Della-Maggiore, 2015; Janssen et al., 2015).

While the above results strongly suggest that contextual information modulates neural markers of motor resonance in M1, the underlying computational process is unclear. Potential, but not mutually exclusive, explanations might relate to (i) a general arousal effect, for example, caused by anticipating an upcoming action (Kilner et al. 2004a; Borroni et al. 2005), (ii) suppression of motor activity due to conflict when contextual and kinematic cues are at odds (Alaerts, Swinnen, & Wenderoth, 2009a; Amoruso et al., 2016; Amoruso & Urgesi, 2016; Senot et al., 2011) or (iii) the integration of kinematic and contextual information according to Bayesian theory as proposed by the predictive coding framework discussed in Kilner, Friston, & Frith, (2007). Here we aim to investigate the latter hypothesis and ask whether M1 combines contextual and kinematic cues according to Bayesian principles. Within this framework, motor resonance can be seen as the result of probabilistic inference in which M1 attempts to infer the most likely cause of an observed action (the posterior) by optimally combining information from observed kinematic data (the likelihood) and the observer’s belief (the contextual prior) according to Bayes’ rule.

To test this hypothesis, we designed two experiments (Figure 1) where we either manipulated the availability of kinematic information (i.e. full-view vs occluded, Fig. 1 top row) or the uncertainty of prior contextual information (i.e. informative vs uninformative contextual cues, Fig. 1 left column) and investigated whether changes in corticomotor excitability reflect the posterior distributions (i.e. the observers belief) as derived from a simple Bayesian model (see Fig. 1 red bars and information in the caption).

The model makes three testable predictions: (i) strongest grip specific motor resonance is observed when kinematic information is available (full-view conditions); (ii) contextual information is sufficient to evoke grip specific motor resonance (occluded condition preceded by informative prior), however, this effect is slightly weaker than when kinematic information is available; and (iii) no grip-specific facilitation when both observed data and prior information are uncertain (occluded condition preceded by uninformative prior).

**Figure 1.**
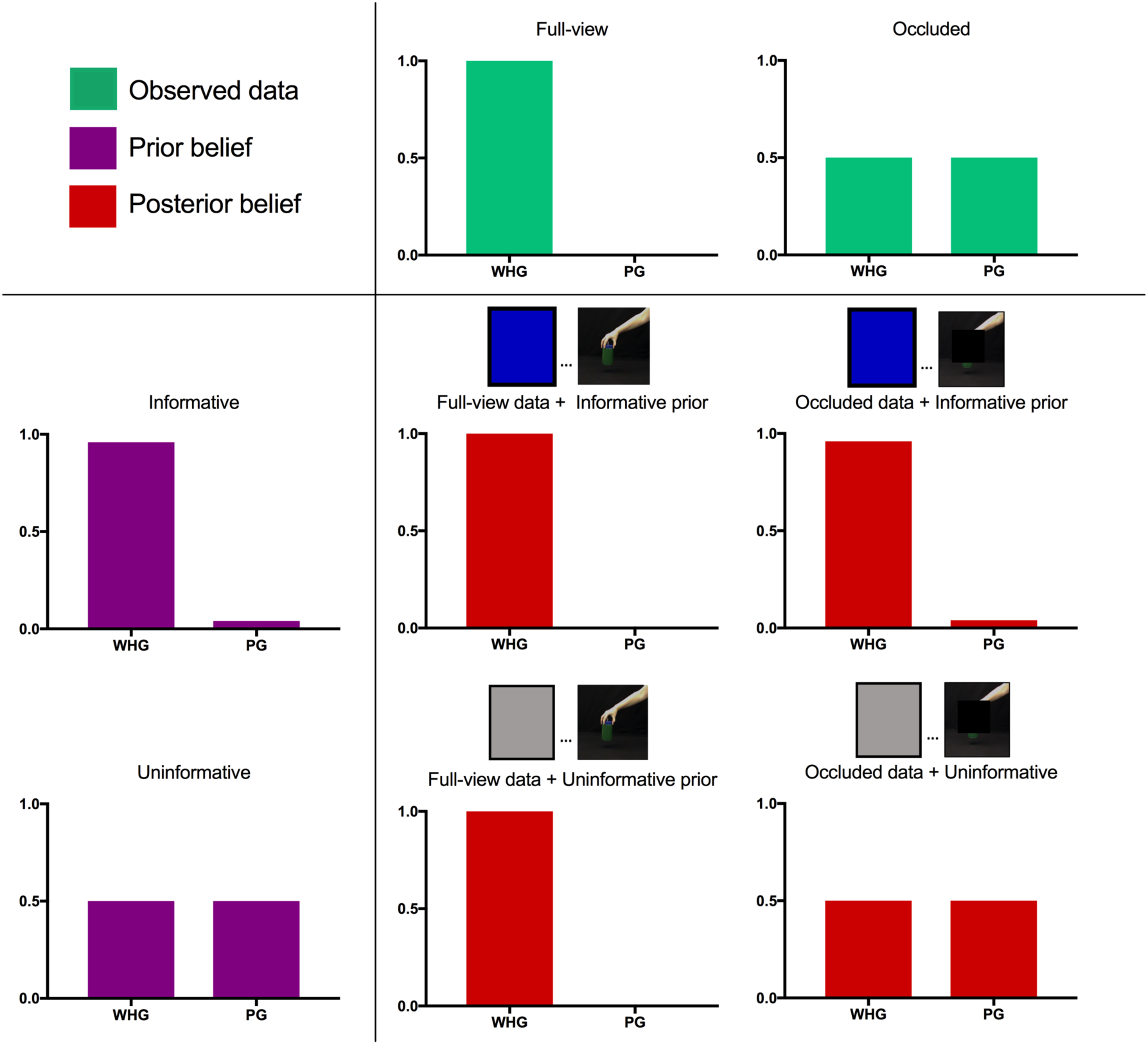
Bayesian model employed to predict changes in corticomotor excitability for different experimental conditions. Bars show grip-specific facilitation in M1 when observing a whole-hand grip (WHG) versus a precision grip (PG). We present a case in which corticospinal facilitation is measured from the abductor digiti minimi (ADM) muscle which has been shown to respond strongly to WHG but not to PG. Videos showing the two grasping actions were preceded by prior information (purple bars in the first column) which was either informative (i.e. in 96% of the trials, blue cues were followed by a WHG) or uninformative (grey cues were followed by WHG in only 50% of the trials). The likelihood function (green bars in the top row) reflects the availability of the kinematic information in the videos i.e. the grasp type was either fully visible (full-view) or the grasp kinematics were covered (occluded). The posterior distributions (red bars) are formed while observing the videos and represent the observer’s belief regarding the grasp type decoded from M1 activity given the observed data (see methods for further details).

## 2. MATERIALS AND METHODS

### 2.1 Bayesian Model

Bayesian models quantify how the brain perceives the external world by combining priors with the incoming sensory information to calculate a posterior distribution: *Posterior* ∝ *Likelihood* × *Prior* In our experiments, participants observed grasping movements (whole-hand or precision grasps) which have been shown to evoke grasp specific muscle activity patterns when measuring corticomotor excitability simultaneously in the first dorsal interosseous (FDI) and the abductor digiti minimi (ADM) muscle. We varied the degree of uncertainty associated with the context (i.e. prior belief regarding an expected grasp type) and the observed kinematics (i.e., sensory input; likelihood). For simplicity we focused only on the extreme cases where the prior belief could either strongly support a specific grasp type (informative, i.e. 96% correct) or could be equally associated with both grasps (uninformative). Similarly, the observed kinematics could either clearly indicate the kinematics of a grasp type (full-view) or could be completely uncertain (occluded).

We used a simple Bayesian model to depict the expected grip-specific motor resonance effect when participants observe full-view or occluded whole-hand grasping (WHG) trials preceded by informative or uninformative cues, when corticospinal excitability is measured from the ADM muscle (Fig. 1). In our experiment informative cues predict the upcoming grasp type with a high accuracy (in 96% of the trials blue cues indicate WHG and white cues indicate PG) while the uninformative are neutral (grey cues) and can be equally associated with the two grasp types (50% probability). Further, we modelled the full-view kinematics as fully reliable sensory input (100% probability for the observed grasp type) while the occluded kinematics predicted a specific grasp type at chance level (50% probability). The posterior probability – the observer’s belief that e.g. a WHG is shown in the video – is derived according to Bayes’ rule:

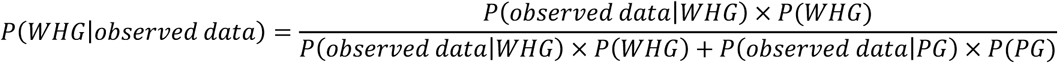

Note that the posterior probabilities for the two grasp types [*P*(*WHG*l*observed data*) and *P*(*PG*l*observed data*)] sum to one.

### 2.2 Participants

24 volunteers participated in Experiment 1 (7 males and 17 females, mean ± SD age 24.5 ± 4.73 years) and 29 in Experiment 2 (13 males and 16 females, mean ± SD age 23.5 ± 2.66 years). All participants were right-handed, as assessed with the Edinburgh Handedness Questionnaire (Oldfield, 1971), naive about the purpose of the experiment and had normal or corrected-to-normal vision. Written informed consent was obtained before the experiments and all subjects were screened for potential contraindications to TMS (Rossi et al., 2009). The experiments were approved by the Kantonale Ethikkommission Zurich and conform to the Declaration of Helsinki (1964).

### 2.3 General Setup

Participants sat on a chair and watched video-clips shown on a computer screen in front of them. Their right hand rested in a neutral position supported by foam pillows (Fig. 2A). In both experiments, subjects observed video-clips showing the right hand of an actor performing grasping actions using either a precision grip (PG) or a whole-hand grip (WHG). Motor-evoked potentials (MEPs) were simultaneously recorded from the First Dorsal Interosseous (FDI) and Abductor Digiti Minimi (ADM) due to their specific differential involvement during the execution and observation of these grip types (Fig. 2B). More specifically, EMG measured during action execution shows that FDI muscle is more active while performing precision grasps and ADM is more active during whole-hand grasps.

**Figure 2.**
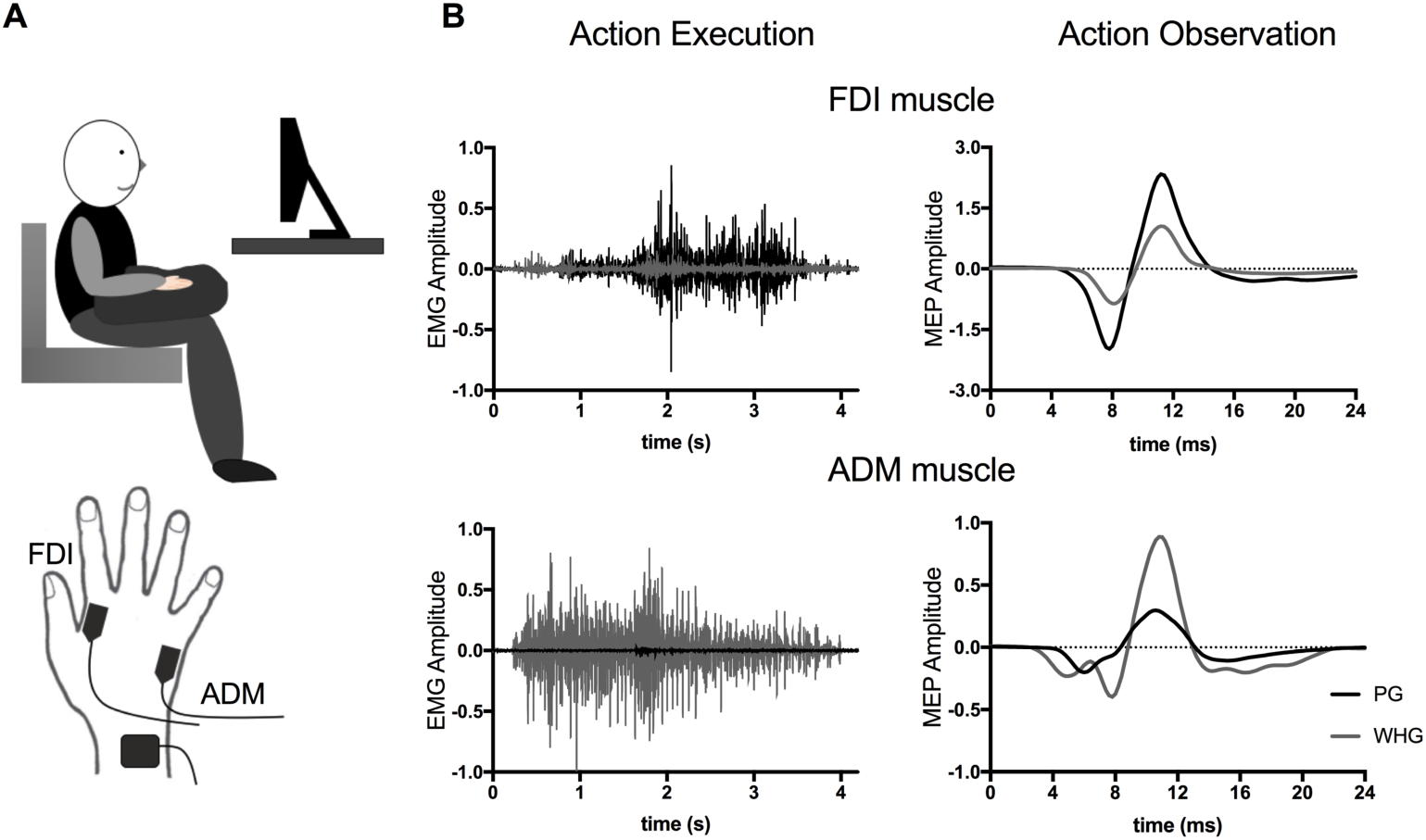
(A) General setup and electrode position (B) Electromyographic (EMG) recordings from the FDI and ADM muscles from a representative subject during the execution and observation of a precision grip (PG, black) and whole-hand grip (WHG, gray). **Action execution:** the subject was instructed to grasp, lift and place a round jar on a stand using either a PG or a WHG. The figure shows the EMG recording from the moment the subject was instructed to start grasping the round jar until the late phase of lifting (before placing the round jar on the stand). **Action Observation:** the subject observed a video showing the hand of an actor reaching for the round jar, grasping it using either a PG or a WHG and lifting it a few centimeters above the surface. Motor-evoked potentials (MEPs) from subject’s right-hand muscles were recorded during the observation of the lifting phase of a movement using a PG and a WHG.

### 2.4 Electromyographic recordings and TMS

Corticomotor excitability was measured using TMS during action observation. MEPs were recorded simultaneously from the First Dorsal Interosseous (FDI) and Abductor Digiti Minimi (ADM) muscles of the right hand using surface electromyography (EMG; Delsys Bagnoli DE-2.1). EMG data were recorded using Signal Software (Version 5.07, Cambridge Electronic Design, UK), sampled at 5000 Hz (CED Power 1401; Cambridge Electronic Design, UK), amplified, band-pass filtered (30–1000 Hz), and stored on a PC for off-line analysis.

TMS was performed by means of a 70 mm figure of eight coil connected to a Magstim200 stimulator (Magstim, Whitland, UK). The coil was positioned over the primary motor cortex (M1) of the left hemisphere, tangentially to the scalp with the handle pointing backwards and laterally at 45° away from the mid-sagittal line, such that the induced current flow was in a posterior anterior direction, i.e., approximately perpendicular to the central sulcus. Since we wanted to make sure that MEPs are consistently elicited in both muscles, we defined the optimal scalp position (hotspot) as the position from which stable responses were simultaneously evoked in both the right FDI and ADM muscles. The resting motor threshold (rMT) was defined as the lowest stimulus intensity evoking MEPs in the right FDI and ADM with an amplitude of at least 50 μV in 5 out of 10 consecutive stimuli (Rossini et al., 2015). In the two experiments, subjects’ rMT, expressed as a percentage of the maximum stimulator output, was on average 48.25 % in Experiment 1 (35 % - 63 %), 46.87 % in Experiment 2 (34 % - 62%). TMS triggering was controlled using Matlab 2013b (The MathWorks, Inc., Natick, USA) in combination with a Teensy 3.1 microcontroller. To reliably record MEPs in both muscles, the stimulation intensity used during both experiments was 130%rMT as described previously (Alaerts et al., 2012; Alaerts, Heremans, Swinnen, & Wenderoth, 2009; de Beukelaar et al., 2016).

Moreover, to ensure a stable coil position during the experiment, the site of stimulation was marked with a Brainsight neuronavigation system (Rogue Research Inc., Montréal, Canada) which allowed online navigation. A Polaris Vicra camera (Northern Digital Inc., Canada) was used to track the coil.

### 2.5 Stimuli and procedure

Stimuli consisted of two video-clips showing the grasping and lifting of a round jar (9 cm diameter, 14.5 cm height, 1.5 kg weight) either with a WHG or PG (Figure 3A). During the WHG all fingers were used to grasp the lid of the jar, whereas for the PG, only the thumb and index finger grasped a small knob mounted on the top of the jar. At the start of the clip the jar was visible in the middle of the screen. Then the right hand of an actor came into view in the upper right corner as a clenched fist. Importantly, up until the opening of the fist the movements were identical in both video-clips. The grip type only became apparent after the initial reaching phase. The jar was grasped and lifted approximately 15 cm above the surface.

In each movie series, we randomized the occurrence of WHG and PG to avoid anticipation of the type of grasp by the observing participants. In half of the video-clips, a PG (n=150) was shown, whereas in the other half, a WHG (n=150) was shown. Furthermore, in order to test whether M1 facilitation is modulated by the reliability of the sensory input, each video-clip was presented either in full-view (n=72 corresponding to 96% accuracy; showing the reach, grasping and lifting kinematics) or occluded (n=72; where the hand-object interaction was occluded by a black square during the grasping and lifting phase). Importantly, in the occluded condition, the black square covered the hand-object interaction after the reaching phase (i.e., participants could not distinguish between grip-types based on kinematic information). The black square remained on the screen until the end of the lift phase. The square occupied approximately 25% of the videos’ area and was sliding with the moving hand (the center of the square was always in the same position as the fingers).

The following procedure was used in both experiments. Participants started the testing session by watching a training block (20 trials) in which they could learn the association between the color cues and the associated grasp-types. They were debriefed at the end of the training block in order to ensure that informative cues were learned and instructions were understood completely.

During the observation of each video-clip, a single TMS pulse was administered either during (i) the reaching phase showing the actor’s hand as a clenched fist (~2 sec after initiation of the video), (ii) the grasp phase when the hand forms according to the grip to be performed (~3.48 sec after initiation) or (iii) the lift phase (~4.92 sec after initiation). When the video-clips were occluded, TMS pulses were sent at the same timepoints as in the full-view condition, corresponding to the reach, grasp and lift phases of the movement.

Furthermore, baseline MEP size was measured by acquiring 20 MEPs while participants watched a black screen. Baseline was tested before the experiment, after performing 10 blocks of the observation task, and at the end of the experiment. Due to technical problems, in experiment 1, the baseline measurements were carried out in 20 out of 24 participants, thus, general effects induced by mirroring will only be analyzed for these subjects.

#### Experiment 1

In order to determine whether M1 grip-specific facilitation is sensitive to the uncertainty of the sensory input, we investigated changes in MEP amplitude during the observation of full-view versus occluded grasping movements. The context or prior knowledge had the same reliability in each trial and was represented by an informative cue which gave participants clear information regarding the type of upcoming grasp action, i.e. before the start of each video-clip, either a blue or a white square was presented to the screen, indicating that the upcoming grasping action would involve a WHG or PG, respectively (Fig.3B upper part).

To ascertain that participants pay attention to the cues and upcoming grip types, they had to indicate after each full-view video-clip whether the cue-grasp video contingency was correct. Participants were informed that the percentage of incorrect cues was rare (4% of all trials) but that it was important to identify them correctly.

Video-clips were divided into 15 blocks, each consisting of a series of 20 grasping movements. For each of the twelve conditions [2 grip types (WHG, PG) × 3 stimulation points (reach, grasp, lift) × 2 video types (full-view, occluded)], 24 MEPs were recorded, resulting in a total of 288 MEPs for each subject and each muscle (FDI, ADM). For the incorrect trials, we only recorded 12 MEPs (6 = WHG and 6 = PG) and this data was excluded from further analyses.

#### Experiment 2

In experiment 2 we aimed to internally replicate experiment 1 while extending the paradigm to a fully factorial design. We investigated whether M1 facilitation is influenced by the reliability of prior knowledge (i.e., informative versus uninformative contextual cues) and sensory input (i.e., full-view versus occluded movement kinematics).

Half of all video-clips were preceded by an informative cue (Fig. 3B upper right part), providing information on the upcoming grip, (i.e., blue square for WHG; white square for PG) while the other half of the video-clips were preceded by an uninformative cue (grey square) which was unspecific and could be followed by either of the two grasps (Fig. 3B lower right part).

**Figure 3.**
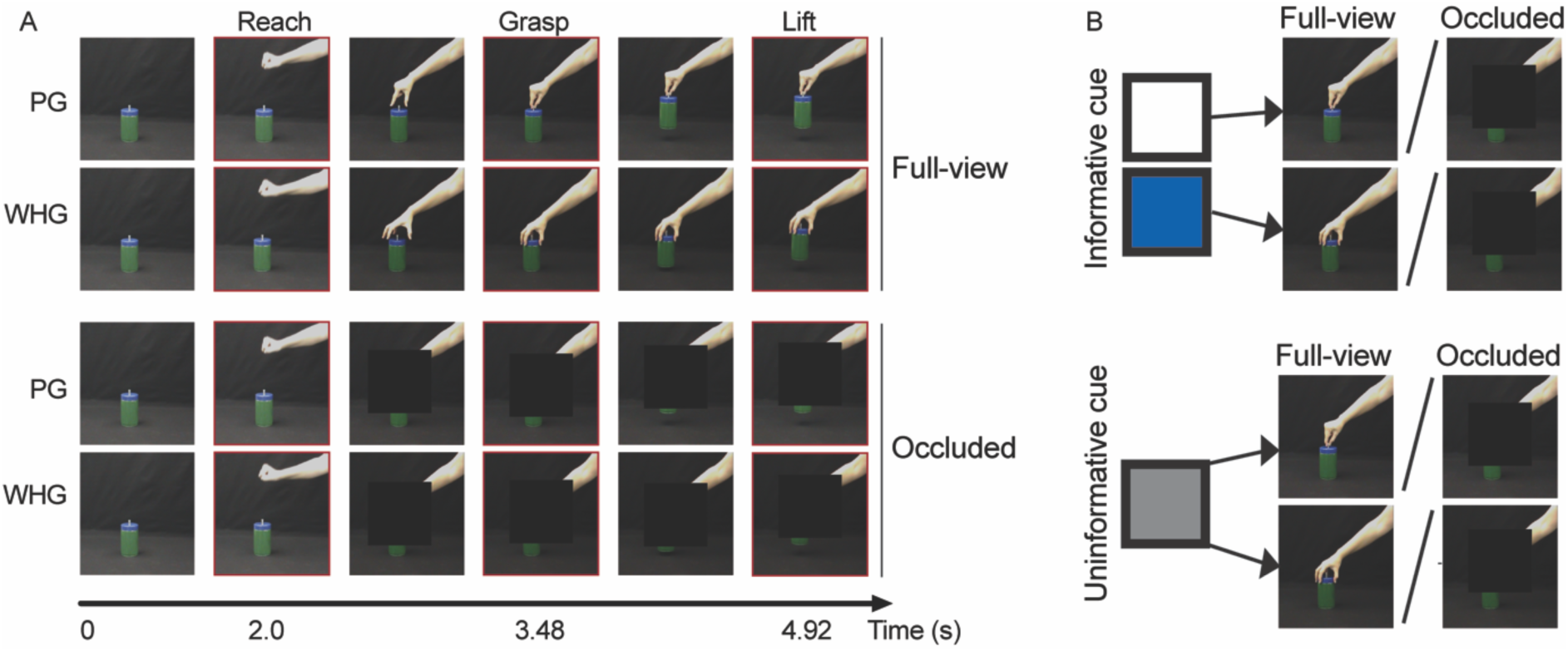
Schematic representation of the videos showing the reaching, grasping and lifting of a round jar by an actor. (A) In both experiments, two types of grasps were presented, a precision grip (PG) or a whole-hand grip (WHG), either in full-view or occluded (i.e., hand-object interaction after the reaching phase was occluded by a black square). During the whole-hand grip, all fingers were used to grasp the lid of the jar while for the precision grip, only the small knob mounted on top of the jar was grasped using the thumb and index finger. Single-pulse TMS was administered at crucial time-points during the unfolding of the movement (depicted by the red boxes surrounding the respective frames): during the reach (2.0 sec), grasp (3.48 sec) and lift (4.92 sec) phases; (B) In experiment 1, each video was preceded by an informative cue: a white square preceded PG videos and a blue square WHG videos (upper part of the figure). In experiment 2, participant observed videos preceded by informative (white, blue) but also uninformative grey cues (i.e., cues which were neutral and could not be used to predict the upcoming grasp type; upper and lower part of the figure).

In order to ensure that participants paid attention to the task they either had to indicate the color of the cue (n=16 participants) or the occluder disappeared at a random time-point before the end of the action (but always after the TMS pulse was sent) and participants had to indicate whether the contingency between the cued and the observed grip type was correct (n=13). This experiment was completed over two sessions that were at least 1 day apart. The procedure was identical in each session.

Subjects were presented with video-clips divided into 15 blocks, each consisting of a series of 20 grasping movements. For each of the twenty-four conditions [2 grip types (WHG, PG) × 3 points (reach, grasp, lift) × 2 video types (full-view, occluded) × 2 cue types (informative, uninformative)], 24 MEPs were recorded across the two sessions, resulting in a total of 576 MEPs for each subject and each muscle. For the incorrect trials, we only recorded 24 MEPs (12 = WHG and 12 = PG) across the two sessions and this data was excluded from further analyses.

### 2.6 Data analysis and statistics

From the EMG data, peak-to-peak amplitudes of the MEPs were calculated using custom-made Matlab scripts (Matlab 2015, Mathworks, Natick, MA, USA). Since background EMG is known to modulate the MEP amplitude, pre-TMS EMG was estimated by determining the root mean square (rms) for a window of 105-5ms before TMS onset. Trials with rms EMG greater than 10 μV were removed from further analysis. Additionally, for each subject, the mean and standard deviation of the background EMG scores were computed and trials with rms EMG larger than the mean + 2.5 SDs were excluded from the analysis. MEPs with a peak-to-peak amplitude smaller than 50 μV were also removed. Following these criteria, a total of 4.34% of all trials were removed in Experiment 1 and 5.25 % in Experiment 2. For the remaining MEPs, the mean peak–to-peak amplitude was calculated separately for each muscle and stimulation condition.

#### General action observation effect on M1

For both experiments, we investigated the non-specific changes in corticospinal excitability during the observation of both grasps compared to baseline. We did this by calculating the difference between the average MEP amplitude during the observation (i.e. collapsing across both grasp types) and the average MEPs acquired during rest in each subject and condition, separately for the FDI and ADM muscles:

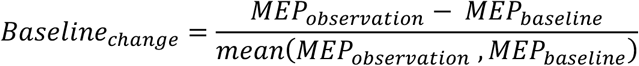

Here, a positive change indicates an overall excitatory effect of observation on corticospinal excitability (interpreted in the literature as proof of a mirror response (Naish et al., 2014)) while a negative change indicates an overall inhibitory effect. We predicted that action observation will produce an increase in corticospinal excitability compared to baseline, therefore, one-sided t-tests were performed for each condition to test specifically whether the baseline change is significantly greater than 0.

#### Grip specific modulation of M1 activity

Second, in accordance to de Beukelaar et al., (2016) we determined whether M1 excitability was modulated in a grip-specific manner, by assessing whether observing a precision versus a whole-hand grip modulated the MEP amplitude:

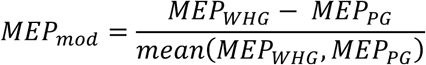

MEP_mod_ was calculated separately for FDI and ADM. Given that both action execution and action observation studies have specified a dissociation between the involvement of these two muscles in each grasp type (e.g., Cavallo, Bucchioni, Castiello, & Becchio, 2013; Lemon et al., 1995; Sartori, Bucchioni, & Castiello, 2012), motor resonance is indicated by positive MEP_mod_ values for the ADM, indicating a stronger facilitation during WHG than PG observation, and negative MEP_mod_ values for the FDI muscle, indicating a stronger facilitation during PG than WHG observation as it can be inferred from figure 2 B. *MEP_mod_* was calculated for each subject, muscle and condition and was subjected to statistical analyses using SPSS 23 (IBM, Armonk, NY, USA).

For **experiment 1**, we employed linear-mixed effects models with *muscle specificity* (FDI, ADM), *video type* (full-view, occluded) and *phase* (reach, grasp, lift) as fixed effects. Subjects were modelled as random effects to account for the high inter-individual variability of MEP measurements. We used a compound symmetry covariance structure, assuming equal variance and equal covariance between the repeated measures.

As a follow up analysis, we performed one sample t-tests to determine whether MEP_mod_ is larger than 0 for ADM and smaller than 0 for FDI as predicted for a grip-specific modulation. Since we had a strong a-priori hypothesis, these tests were one-sided.

The significance threshold alpha=.05 was chosen for all statistical tests and FDR corrections (Benjamini & Hochberg, 1995) for multiple comparisons were applied when necessary. Results are reported in the figures as mean ± standard error of the mean (SEM).

In addition to classical analyses based on p-values to reject the null-hypothesis, we employed Bayesian repeated-measures ANOVAs using JASP software (Version 0.8.1.0, https://jasp-stats.org/). These tests produce a Bayes factor (BF10) which is a graded measurement indicating evidence provided by the data in favor of the null (H_0_) or the alternative hypothesis (H_1_) (Apšvalka, Ramsey, & Cross, 2018; Dienes, 2016; Dienes & Mclatchie, 2017; Rouder, Speckman, Sun, Morey, & Iverson, 2009). In detail, BF10=1 demonstrates that the data does not favor either hypothesis while values > 1 suggest increasing evidence for the alternative hypothesis (Kass & Raftery, 1995; Rouder, Morey, Verhagen, Swagman, & Wagenmakers, 2017). In general, a BF_10_ > 10 is considered strong evidence for the alternative hypothesis while BF_10_ < 1 indicates evidence in favor of the null hypothesis (adapted from Jeffreys, 1961).

Moreover, we computed the BF_inclusion_ factor across matched models which compares the evidence for all the models containing an effect of interest versus the models not containing it. Since increasing the number of factors will automatically increase the number of models to be compared in an analysis, we used the BF_inclusion_ to facilitate the interpretations and decide which models are more plausible and better supported by the data (Wagenmakers et al., 2017). The Bayesian repeated-measures ANOVA contained the same factors (i.e., *muscle specificity*, *video* and *phase*) as the mixed effects models.

Analyses of **experiment 2** were analogous to Experiment 1. Importantly, separate linear mixed-effects models were applied to MEP_mod_ data obtained with informative cues and to data obtained with non-informative cues. Each of these statistical models contained the factors *muscle specificity* (FDI, ADM), *video type* (full-view, occluded) and *phase* (reach, grasp, lift) were employed.

#### Analysis of combined data from experiment 1 and 2 for the full-view Informative Cue Condition

Since participants observed identical full-view videos preceded by informative cues in both experiments, we combined the results from experiment 1 and 2 in order to enlarge the sample size to n=53 (Experiment 1 = 24; Experiment 2 = 29). These data were subjected to a *video type* × *muscle specificity* × *phase* (reach, grasp, lift) mixed effects model and t-tests were employed to determine whether MEP_mod_ differed from 0 as predicted for a grip-specific modulation. We were particularly interested in testing the hypothesis that motor resonance is slightly stronger for contextual + kinematic than for contextual information only. Therefore, we performed a separate *video type* × *muscle specificity* mixed effects model including only the grasp and the lift phase, since only these phases provided kinematic information regarding the grip type in the full-view videos.

## 3. RESULTS

### Experiment 1: Observation of full-view and occluded movements preceded by informative cues

#### General changes in excitability during movement observation

We first tested whether movement observation in general (i.e. collapsed over the different grasp types) influences corticomotor excitability. For both muscles, corticomotor excitability was larger during action observation than during baseline measurements (Supporting Information table S1). One-sided t tests revealed that changes from baseline measured in the FDI muscle were significantly larger than zero in all conditions (p_FDR_<=.01), indicating a general ‘mirroring’ response. For the ADM muscle, the changes from baseline were also larger than zero, but the differences were smaller and did not reach significance after correction for multiple comparisons.

#### Grip-specific changes in excitability during movement observation

M1 was modulated in a grip-specific manner (i.e., on average we found MEP_mod_ < 0 for FDI and MEP_mod_ > 0 for ADM) during the observation of both full-view and occluded actions, as it can be seen in Fig. 4.

**Figure 4.**
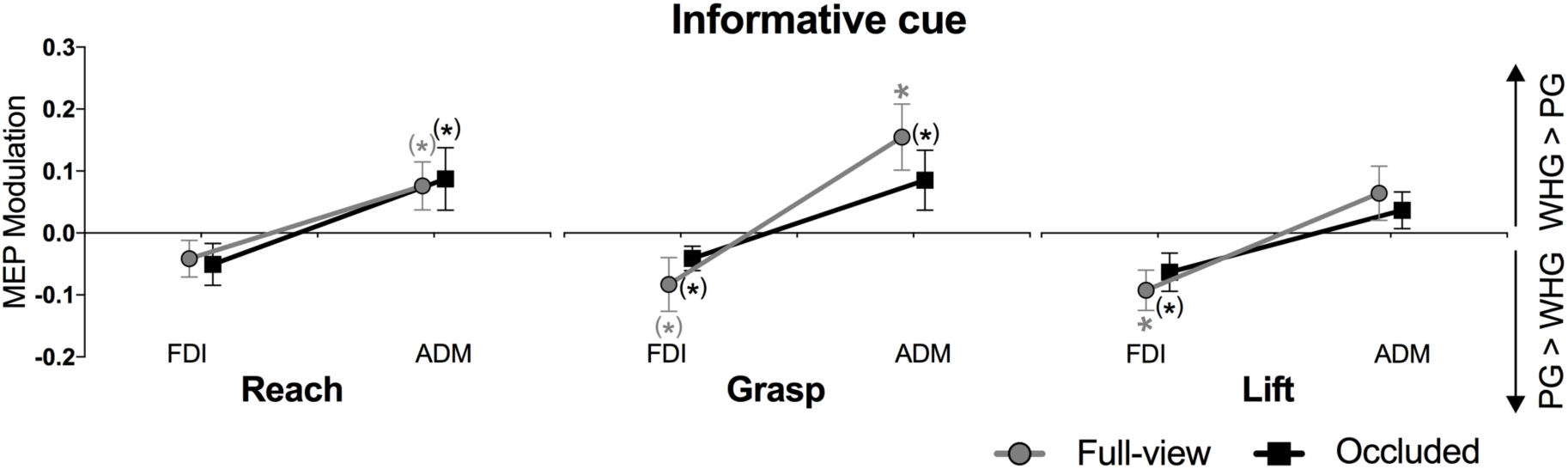
Results of Experiment 1: MEP modulation (MEP_mod_) during the observation of full-view (gray lines) or occluded (black lines) videos depicting a whole-hand grip (WHG) or a precision grip (PG); Results are presented separately for each movement phase (reach, grasp, lift) and muscle (FDI, ADM); Values > 0 indicate higher facilitation during WHG than PG observation, while values < 0 indicate increased facilitation during PG than WHG observation; ∗: p<.05 (FDR corrected one-sided t tests),(∗): p<.05 (uncorrected);

Both the mixed-effects analyses as well as Bayesian statistics yielded strong evidence for a main effect of muscle specificity (table 1; p<.0001, BF_inclusion_=8.1*10^7^) but no other main effects or interactions reached significance (all p>.2, BF_inclusion_<.31). The same result pattern was obtained when the occluded and the full-view videos were analyzed with separate mixed-effects models (table 2, main effect of muscle specificity; *occluded* p<.001, BF_inclusion_=524; *full-view* p<.001, BF_inclusion_=6.7*10^4^). This indicates that corticomotor excitability of the two muscles was differentially modulated by the cued grasp type irrespective of whether kinematics were occluded or not and irrespective of the phase. Note that the grip type was not yet visible during the reach phase (where the hand was still forming a fist). This explains why muscle specific changes in this phase were overall smaller than during the grasp phase even though this effect did not reach significance (muscle specificity × phase interaction: p=.52 and also the Bayesian RM-ANOVA provided evidence against a muscle × phase effect: BF_inclusion_=.105).

**Table 1.** Experiment 1 (N=24): linear mixed effects and Bayesian Repeated-measures ANOVA

|  | Mixed-effects analyses |  | Bayesian RM - ANOVA |  |
| --- | --- | --- | --- | --- |
|  | F values | p-values | BF<sub>10</sub> | BF<sub>inclusion</sub> |
| *Main effects* |  |  |  |  |
| **Muscle (FDI, ADM)** | **42.77** | **.0001** | **7.7350\*10<sup>7</sup>** | **8.1\*10<sup>7</sup>** |
| Video (full-view, occluded) | 0.039 | .859 | 0.129 | 0.136 |
| Phase (reach, grasp, lift) | 1.295 | .276 | 0.108 | 0.147 |
| *Interactions* |  |  |  |  |
| Muscle x Phase | 0.655 | .521 | 1.2*10<sup>6</sup> | 0.105 |
| Muscle x Video | 1.222 | .27 | 3.317*10<sup>6</sup> | 0.308 |
| Phase x Video | 0.048 | .954 | 9.879*10<sup>-4</sup> | 0.074 |
| Muscle x Phase x Video | 0.733 | .481 | 7.64*10<sup>2</sup> | 0.202 |
| Degrees of freedom = 253 |  |  |  |  |
| Significant p-values represented in bold (p<0.05) |  |  |  |  |

**Table 2.** Experiment 1 (N=24): linear mixed effects and Bayesian Repeated-measures ANOVA

| **Table 2** Experiment 1 (N=24): linear mixed effects and Bayesian Repeated-measures ANOVA results |  |  |  |  |
| --- | --- | --- | --- | --- |
|  | **Mixed-effects analyses** |  | **Bayesian RM - ANOVA** |  |
|  | F values | p-values | BF<sub>10</sub> | BF<sub>inclusion</sub> |
| Full-view condition |  |  |  |  |
| *Main effects* |  |  |  |  |
| **Muscle (FDI, ADM)** | **28.895** | **.0001** | **6.67\*10<sup>4</sup>** | **6.76\*10<sup>4</sup>** |
| Phase (reach, grasp, lift) | 0.837 | .436 | 0.125 | 0.14 |
| *Interactions* |  |  |  |  |
| Muscle x Phase | 1.247 | .291 | 3.105*10<sup>3</sup> | 0.331 |
| Occluded condition |  |  |  |  |
| *Main effects* |  |  |  |  |
| **Muscle (FDI, ADM)** | **16.32** | **.0001** | **529.97** | **524.1** |
| Phase (reach, grasp, lift) | 0.548 | .58 | 0.119 | 0.106 |
| *Interactions* |  |  |  |  |
| Muscle x Phase | 0.14 | .87 | 6.965 | 0.124 |
| Degrees of freedom = 115 |  |  |  |  |
| Significant p-values represented in bold (p<0.05) |  |  |  |  |

Follow-up analyses testing whether MEPmod deviated significantly from zero revealed several trends towards significance (particularly for the grasp phase), however, only grip-specific facilitation of ADM during the grasp phase of full-view movements (t=2.89; P_FDR_ = .004) and grip-specific facilitation of FDI during the lift phase of full-view movements (t=-2.82; P_FDR_=.005) survived correction for multiple comparisons (table 3). In summary, this pattern of MEP modulation indicates that contextual cues can activate motor representations of different grip types in M1 even in the absence of kinematic stimuli.

**Table 3.** Experiment 1 (N=24): Descriptive statistics and results of one-sided t tests performed with the MEP_mod_.

|  | Full-view FDI |  |  | Full-view ADM |  |  | Occluded FDI |  |  | Occluded ADM |  |  |
| --- | --- | --- | --- | --- | --- | --- | --- | --- | --- | --- | --- | --- |
|  | Reach | Grasp | Lift | Reach | Grasp | Lift | Reach | Grasp | Lift | Reach | Grasp | Lift |
| mean | -0.04 | -0.08 | -0.09 | 0.08 | 0.15 | 0.06 | -0.05 | -0.04 | -0.06 | 0.09 | 0.08 | 0.04 |
| SD | 0.14 | 0.21 | 0.16 | 0.19 | 0.26 | 0.21 | 0.16 | 0.09 | 0.15 | 0.25 | 0.24 | 0.14 |
| p-values | .088 | .034 | **.005** | .031 | **.004** | .077 | .074 | .023 | .026 | .049 | .045 | .112 |

The table shows the grip-specific motor resonance which is indicated by values >0 in the ADM muscle (i.e., a stronger facilitation during WHG than PG observation) and values <0 for the FDI muscle. p-values that survived multiple comparisons correction are represented in bold.

### Experiment 2: Observation of full-view and occluded movements preceded by informative and uninformative cues

#### General changes in excitability during movement observation

Changes from baseline measured in both muscles were significantly larger than 0 for all the experimental conditions (all FDR corrected one-sided t tests p<=.002; Supporting Information table S2). This suggests that the observation of a movement or simply knowledge about the presence of a movement can lead to increased corticospinal excitability compared to baseline.

#### Grip-specific changes in excitability during movement observation

M1 was modulated in a grip-specific manner (average MEP_mod_ for FDI were smaller than 0 and for ADM larger than 0) during the observation of both full-view and occluded actions when an informative cue was presented before the start of the action (Figure 5 A). Accordingly, mixed-effects analyses and Bayesian statistics (table 4) revealed a significant *muscle specificity* main effect (p <.001, BF_inclusion_=6.21*10^15^) thus replicating experiment 1. Additionally, similar to the previous experiment, the muscle specific changes in the reach phase were overall smaller than during the grasp and lift phase, which was confirmed by a significant muscle specificity × phase interaction (p <.001, BF_inclusion_=8.73; all other effects p>.078, BF_inclusion_ <.83). By contrast, when uninformative cues were used, evidence for motor resonance (i.e. a muscle specific modulation of MEP_mod_) was only observed when the kinematics were visible during the Grasp and Lift phase (Figure 5B, green symbols) as indicated by a significant *muscle specificity* × *video type* interaction (p<.0001, BF_inclusion_=2.08*10^4^). This is an important control result, indicating that even though corticomotor excitability was generally increased for all conditions (see above), grip specific modulation was absent when participants received neither kinematic nor contextual information.

**Figure 5.**
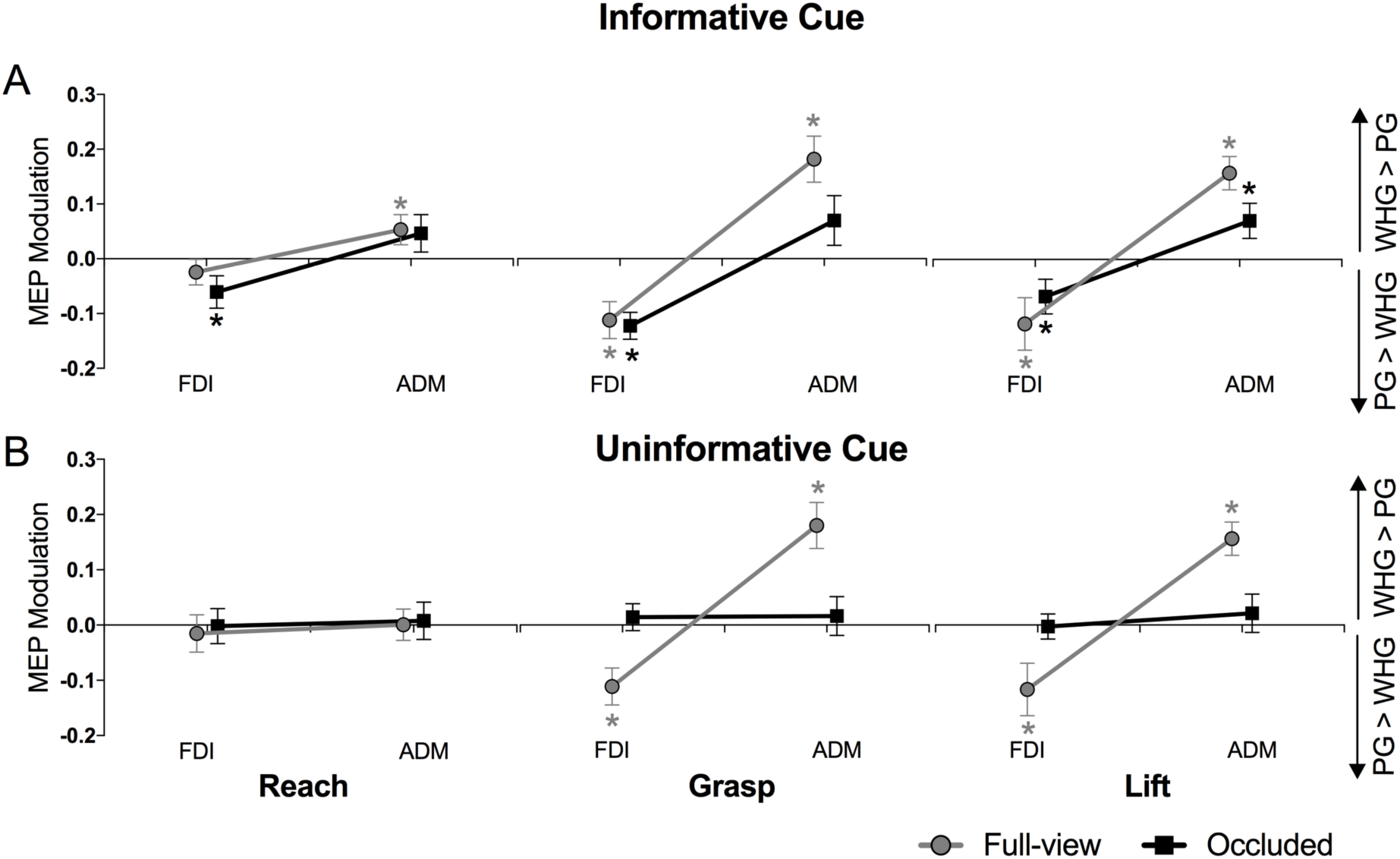
Results of experiment 2. MEP modulation (MEP_mod_) during the observation of full-view (gray lines) or occluded (black lines) videos depicting a whole-hand grip (WHG) or a precision grip (PG); **A:** Videos were preceded by informative cues; **B:** Videos were preceded by uninformative cues; Results are presented separately for each movement phase (reach, grasp, lift) and muscle (FDI, ADM); Values > 0 indicate higher facilitation during WHG than PG observation, while values < 0 indicate increased facilitation during PG than WHG observation; ∗: p<.05 (FDR corrected one-sided t tests);

**Table 4.** Experiment 2 (N=29): Linear mixed effects and Bayesian Repeated-measures ANOVA results

|  | Mixed-effects analyses |  | Bayesian RM - ANOVA |  |
| --- | --- | --- | --- | --- |
|  | F values | p-values | BF<sub>10</sub> | BF<sub>inclusion</sub> |
| Informative cue |  |  |  |  |
| *Main effects* |  |  |  |  |
| **Muscle (FDI, ADM)** | **89.85** | **.0001** | **5.51\*10<sup>15</sup>** | **6.21\*10<sup>15</sup>** |
| Phase (reach, grasp, lift) | 0.056 | .946 | 0.034 | 0.032 |
| Video (full-view, occluded) | 3.129 | .078 | 0.386 | 0.562 |
| *Interactions* |  |  |  |  |
| **Muscle x Phase** | **5.69** | **.004** | **1.531\*10<sup>15</sup>** | **8.726** |
| Muscle x Video | 3.34 | .069 | 2.626*10<sup>15</sup> | 0.824 |
| Phase x Video | 0.519 | .595 | .001 | 0.098 |
| Muscle x Phase x Video | 1.763 | .173 | 2.79*10<sup>13</sup> | 0.391 |
| Uninformative cue |  |  |  |  |
| *Main effects* |  |  |  |  |
| **Muscle (FDI, ADM)** | **27.171** | **.0001** | **1.07\*10<sup>4</sup>** | **1.07\*10<sup>4</sup>** |
| Phase (reach, grasp, lift) | 0.429 | .652 | 0.045 | 0.045 |
| Video (full-view, occluded) | 0.464 | .496 | 0.143 | 0.152 |
| *Interactions* |  |  |  |  |
| **Muscle x Phase** | **4.783** | **.009** | **1.74\*10<sup>3</sup>** | **4.662** |
| **Muscle x Video** | **24.905** | **.0001** | **3.245\*10<sup>7</sup>** | **2.08\*10<sup>4</sup>** |
| Phase x Video | 0.482 | .618 | .001 | 0.086 |
| **Muscle x Phase x Video** | **6.398** | **.002** | **1.704\*10<sup>7</sup>** | **28.868** |
| Degrees of freedom = 308 |  |  |  |  |
| Significant p-values represented in bold (p<0.05) |  |  |  |  |

One-sided t-tests (table 5) revealed that observing full-view grasping kinematics significantly modulated the grip-specific facilitation in the FDI and ADM muscles (p_FDR_<=.01). When movement kinematics were occluded but preceded by informative cues, the grip-specific facilitation followed the same pattern and reached significance in all but one condition (p_FDR_<=.02; ADM occluded grasp p=.068). At the start of the observed movements (i.e., during the observation of the clenched fist) the grip-specific modulation was significant both FDI and ADM muscles but only in one of the conditions (FDI occluded p_FDR_=.02; ADM full-view p_FDR_=.03). As expected, this modulation was completely abolished when the cue was uninformative and no kinematic information was provided (all p-values for the occluded neutral cue conditions >=.27).

**Table 5.** Experiment 2 (N=29): Descriptive statistics and results of one-sided t tests performed with the MEPmod.

| Informative cue |  |  |  |  |  |  |  |  |  |  |  |  |
| --- | --- | --- | --- | --- | --- | --- | --- | --- | --- | --- | --- | --- |
|  | *Full-view FDI* |  |  | *Full-view ADM* |  |  | *Occluded FDI* |  |  | *Occluded ADM* |  |  |
|  | Reach | Grasp | Lift | Reach | Grasp | Lift | Reach | Grasp | Lift | Reach | Grasp | Lift |
| mean | -0.02 | -0.11 | -0.12 | 0.05 | 0.18 | 0.16 | -0.06 | -0.12 | -0.07 | 0.05 | 0.07 | 0.07 |
| SD | 0.13 | 0.18 | 0.26 | 0.15 | 0.23 | 0.16 | 0.16 | 0.13 | 0.17 | 0.18 | 0.24 | 0.17 |
| p-value | .156 | **<.01** | **.010** | **.032** | **<.01** | **<.01** | **.025** | **<.01** | **.021** | .094 | .068 | **.018** |
| Uninformative cue |  |  |  |  |  |  |  |  |  |  |  |  |
|  | *Full-view FDI* |  |  | *Full-view ADM* |  |  | *Occluded FDI* |  |  | *Occluded ADM* |  |  |
|  | Reach | Grasp | Lift | Reach | Grasp | Lift | Reach | Grasp | Lift | Reach | Grasp | Lift |
| mean | -0.015 | -0.11 | -0.10 | .0006 | 0.18 | 0.16 | -0.002 | 0.014 | -0.003 | 0.008 | -0.02 | 0.021 |
| SD | 0.18 | 0.19 | 0.18 | 0.15 | 0.19 | 0.18 | 0.17 | 0.13 | 0.12 | 0.18 | 0.19 | 0.19 |
| p-value | .326 | **<.01** | **<.01** | .491 | **<.01** | **<.01** | .474 | .283 | .451 | .409 | .322 | .271 |

The table shows the grip-specific motor resonance which is indicated by values $>0$ in the ADM muscle (i.e., a stronger facilitation during WHG than PG observation) and values $<0$ for the FDI muscle. p-values that survived multiple comparisons correction are represented in bold.

#### Experiment 1+2

Observation of full-view and occluded movements preceded by informative cues Experiment 1 and Experiment 2 revealed that an informative cue is sufficient to evoke grasp specific modulation of MEP amplitudes even in the absence of visible kinematics (Fig. 4 and 5A). This modulatory effect tended to be smaller for the occluded (black symbols) than for the full-view condition (green symbols), however, statistics failed to reveal significant effects potentially because statistical power was low. Here we pooled the data across experiments (Figure 6; N=53) and found a significant muscle × video type interaction (p=.037) and a significant muscle × phase interaction (p=.01), in addition to the main effect of muscle (p<.001). The Bayesian RM-ANOVA only revealed evidence in favor for the muscle specificity model (table 6; BF_inclusion_=8.1*10^7^).

**Figure 6.**
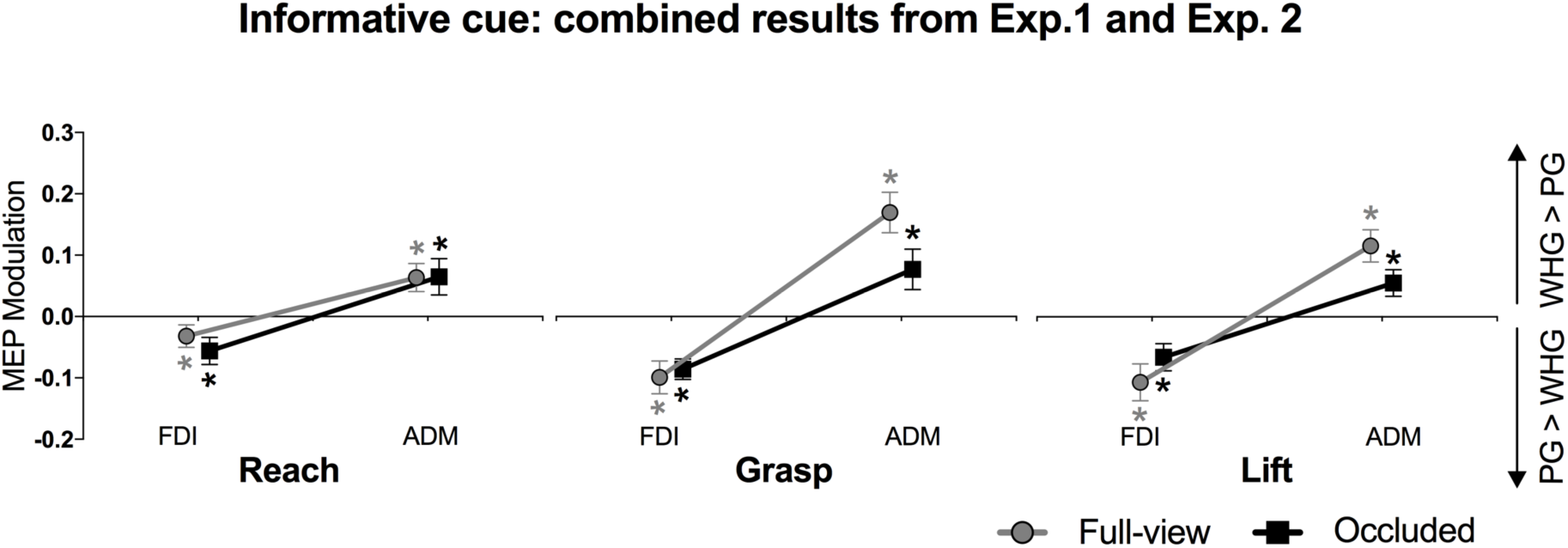
Combined results from Experiments 1 and 2 for the informative cue condition. Significant grip specific facilitation was observed in both muscles (FDI, ADM) in all the phases of the movement. ∗: p<.05 (FDR corrected one-sided t tests);

**Table 6.** Experiment 1+2 (N=53): Linear mixed effects and Bayesian Repeated-measures ANOVA results

|  | Mixed-effects analyses |  | Bayesian RM - ANOVA |  |
| --- | --- | --- | --- | --- |
| | F values | p-value | $BF_{10}$ | $BF_{inclusion}$ |
| *Main effects* |  |  |  |  |
| **Muscle (FDI, ADM)** | **129.637** | **.0001** | **$7.735*10^7$** | **$8.1*10^7$** |
| Video (full-view, occluded) | 1.953 | .163 | 0.129 | 0.136 |
| Phase (reach, grasp, lift) | 0.402 | .669 | 0.108 | 0.147 |
| *Interactions* |  |  |  |  |
| **Muscle x Phase** | **4.618** | **.01** | **1.2\*10<sup>6</sup>** | **0.105** |
| **Muscle x Video** | **4.354** | **.037** | **3.317\*10<sup>6</sup>** | **0.308** |
| Phase x Video | 0.45 | .638 | 9.879*10<sup>-4</sup> | 0.074 |
| Muscle x Phase x Video | 2.192 | .113 | 7.64*10<sup>2</sup> | 0.202 |
| Degrees of freedom = 572 |  |  |  |  |
| Significant p-values represented in bold (p<0.05) |  |  |  |  |

In order to better understand whether cue + kinematic information causes stronger motor resonance than cue information alone we calculated a reduced statistical model including only the lift and grasp phase, i.e. those phases where the kinematics clearly differ between grasps and one would expect a differential effect for full-view versus occluded videos. Both the mixed effects model and the Bayesian RM ANOVA (table 7) revealed a significant muscle specificity × video effect (p=.005, BF_inclusion_=7.17) indicating that cue + kinematic information causes a significantly stronger change in MEP_mod_ than cue information alone. However, this effect is small, and effect sizes differ only slightly between observing the videos in the the cue only (dgrasp=.66 and d_lift_=.53) and the cue + full-view kinematics condition (d_grasp_=.88 and d_lift_=.77).

**Table 7.**
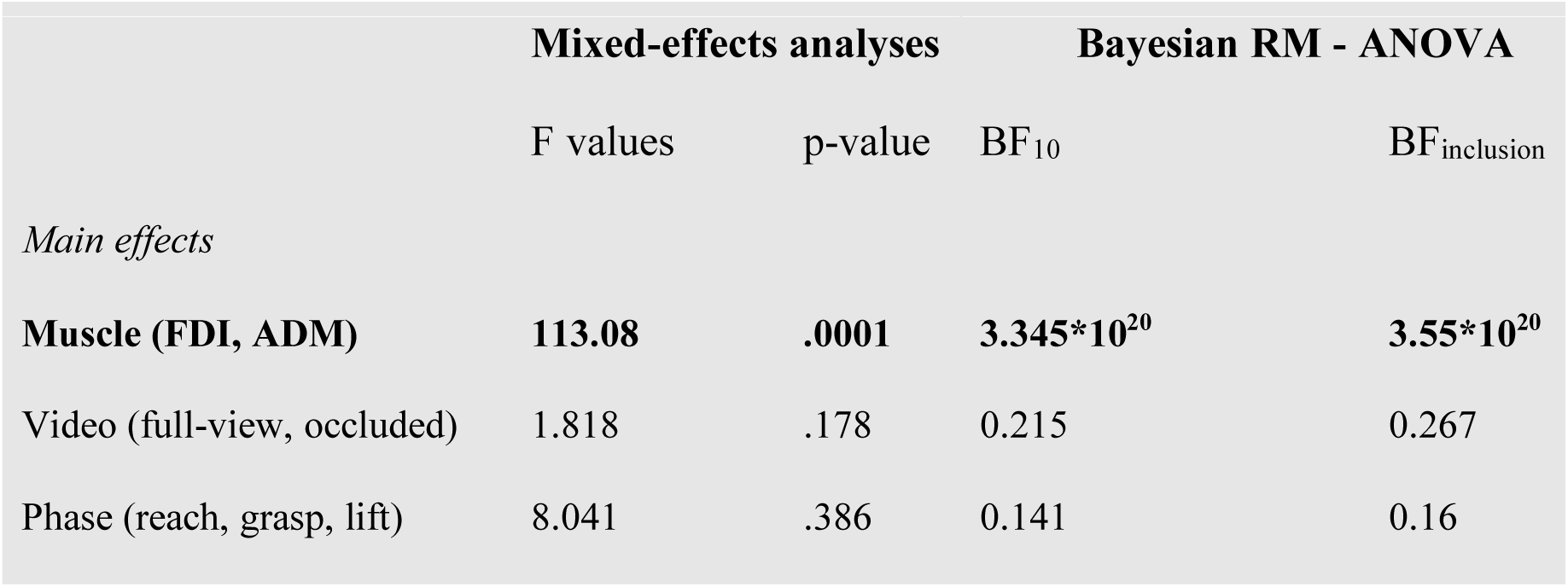
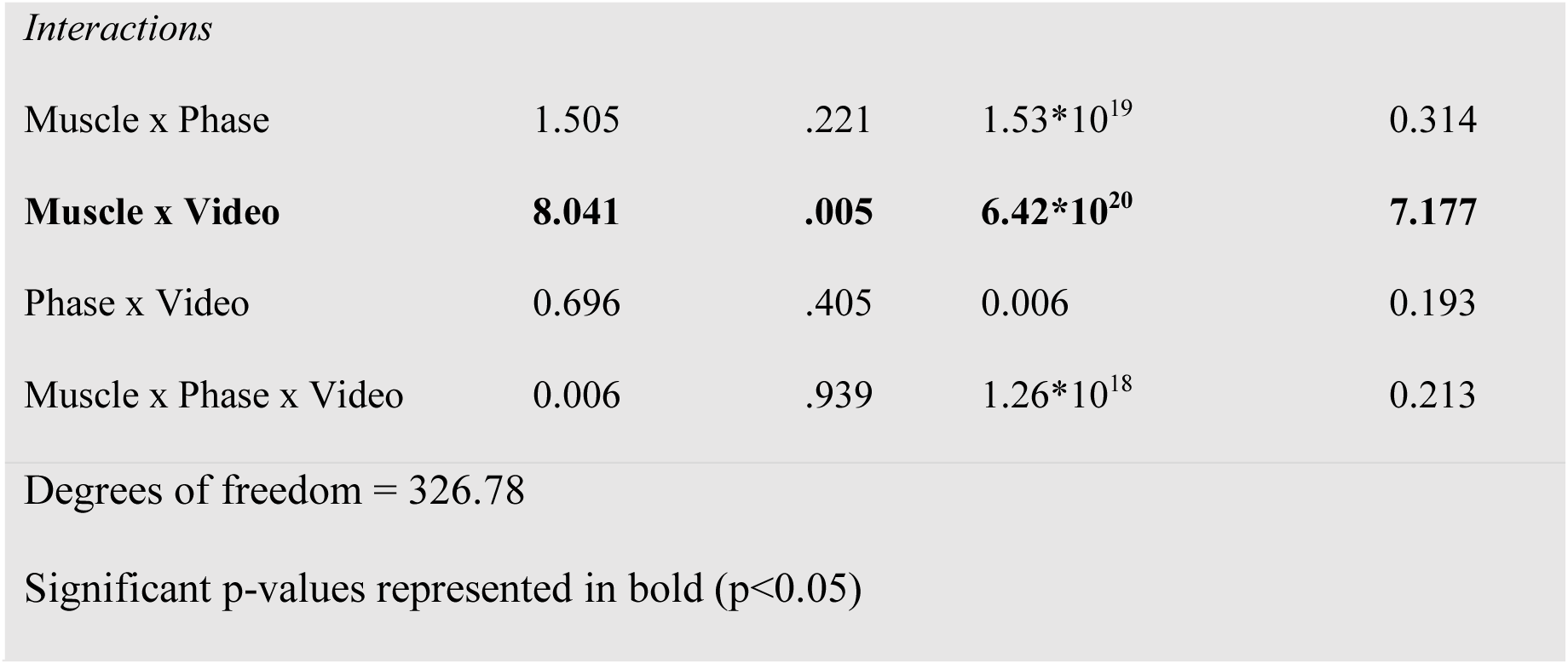
Experiment 1+2 (N=53): Reduced linear mixed effects and Bayesian Repeated-measures ANOVA models (grasp and lift phase)

|  | Mixed-effects analyses |  | Bayesian RM - ANOVA |  |
| --- | --- | --- | --- | --- |
| | F values | p-value | $BF_{10}$ | $BF_{inclusion}$ |
| *Main effects* |  |  |  |  |
| **Muscle (FDI, ADM)** | **113.08** | **.0001** | **3.345\*10<sup>20</sup>** | **3.55\*10<sup>20</sup>** |
| Video (full-view, occluded) | 1.818 | .178 | 0.215 | 0.267 |
| Phase (reach, grasp, lift) | 8.041 | .386 | 0.141 | 0.16 |

| *Interactions* |  |  |  |  |
| --- | --- | --- | --- | --- |
| Muscle x Phase | 1.505 | .221 | $1.53*10^{19}$ | 0.314 |
| **Muscle x Video** | **8.041** | **.005** | **$6.42*10^{20}$** | **7.177** |
| Phase x Video | 0.696 | .405 | 0.006 | 0.193 |
| Muscle x Phase x Video | 0.006 | .939 | $1.26*10^{18}$ | 0.213 |
| Degrees of freedom = 326.78 |  |  |  |  |
| Significant p-values represented in bold ($p<0.05$) | | | | |

## 4. DISCUSSION

In the present study we investigated whether motor resonance evoked by observing movement kinematics and/or contextual cues can be understood within a Bayesian framework. We operationalized motor resonance as muscle specific changes in corticomotor excitability when observing different grasping actions and systematically varied the uncertainty of kinematic versus contextual information. Our data confirmed three major predictions that were derived from a simple Bayesian model:

First, observing the kinematics of a precision versus whole hand grip significantly modulated corticospinal excitability of FDI and ADM according to a grip specific muscle activity pattern. This indicates that kinematic information causes strong motor resonance (Cohen’s d=.88 for Exp. 1 and 2) via bottom-up mechanisms linking visual motion stimuli to motor representations. This effect was observed irrespective of whether additional contextual information was available or not (Exp. 2). Second, contextual information was sufficient to evoke significant grip specific motor resonance even when kinematic information was absent as in the occluded videos or ambiguous as during the reach phase. This suggests that contextual cues activate top-down mechanisms which, in turn, drive motor representation in M1. Importantly, muscle specific changes of MEP_mod_ were slightly but significantly smaller when evoked by contextual information only (which was less certain since it was correct for only 96% of the trials) than when kinematic information was available (Exp. 1 and 2; Cohen’s d occluded >=.54; Cohen’s d full-view>=.77).

Finally, there was no grip-specific facilitation when neither kinematic nor context information was available (i.e. for uninformative cues and occluded videos/reach phase) which served as a control condition indicating that changes in corticomotor excitability reflect grip specific motor resonance rather than general arousal effects.

An important contribution of our design over previous human action observation studies was to employ a task in which the initial kinematics (i.e., reaching) were the same although the final kinematics (i.e., grasp and lift) were different. Participants observed the start of the movements but had no way of knowing which grasp type was hidden behind the occluder – unless informative contextual cues were present. This allowed us to demonstrate that grip-specific facilitation of the human corticospinal system is present when the hand-object interaction is hidden from view indicating that it can be driven by bottom-up kinematic and top-down contextual information. Similar modulation was observed by Umiltà and colleagues, (2001) in the premotor area F5 of macaque monkeys - where a subset of premotor mirror neurons were found to be active during the observation of partially occluded movements. This activation was considered to be important for understanding the goal of observed motor acts, however, it is still unclear which mechanism is reflected by this activity in F5.

### General versus grip specific changes of corticomotor excitability during action observation

Particularly for experiment 2, we demonstrated a general significant facilitation of corticospinal excitability during movement observation, i.e. MEPs elicited during grasp observation were larger than during baseline measurements which is in line with previous studies (Alaerts et al., 2009; Alaerts, Van Aggelpoel, Swinnen, & Wenderoth, 2009; Fadiga, Fogassi, Pavesi, & Rizzolatti, 1995; Gangitano, Mottaghy, & Pascual-Leone, 2001; Hardwick, Mcallister, Holmes, & Edwards, 2012; Hogeveen & Obhi, 2012; Kaneko, Yasojima, & Kizuka, 2007; Lepage, Tremblay, & Théoret, 2010; Strafella & Paus, 2000). Note, however, that this general increase in corticomotor excitability provides different information than grip specific facilitation of the FDI versus ADM muscle. First, both measurements are mathematically orthogonal, i.e. the general facilitation is calculated as a change in average MEP amplitude (collapsed across the different grip types) while MEP_mod_ reflects the grip specific modulation of MEP amplitude over and above these average amplitude changes. EMG measurements as well as previous research investigating the observation of different grip types have showed that FDI contributes to both WHG and PG whereas ADM is specifically involved in the formation of WHG (Bunday, Lemon, Kilner, Davare, & Orban, 2016; Davare, Lemon, & Olivier, 2008; Davare, Montague, Olivier, Rothwell, & Lemon, 2009; Sartori, Betti, & Castiello, 2013). Accordingly, the general MEP facilitation is more pronounced for FDI than for ADM (as found in Exp. 1) while MEP_mod_ seems to be slightly larger for ADM than for FDI (see also: Kilner et al. 2004; Alaerts et al. 2012; de Beukelaar et al. 2016; Jacquet et al. 2016).

Second, the results of experiment 2 revealed a significant increase of general MEP amplitude even when neither contextual nor kinematic cues were available while MEP_mod_ did not change, indicating the absence of any grip specific motor resonance effect. This is in line with previous work reporting unspecific facilitation of the motor system within an action observation context (Grafton, Fadiga, Arbib, & Rizzolatti, 1997; Murata et al., 1997; Valchev et al., 2015), however, it is likely that these changes are due to widespread effects of attention or arousal. As pointed out by Cavallo and colleagues (2014), illusory ‘mirror’ responses might be observed when muscle-specific designs (i.e., two action/two-muscle designs) are not used. This is an important aspect when describing the functional role of motor resonance for action understanding. Analyzing general changes in corticospinal excitability doesn’t offer any specific information regarding the mirrored kinematics used in the observed movements. We could speculate that when both contextual cues and movement kinematics were ambiguous in Exp. 2, the increase in MEP amplitude that we observed could indicate that, compared to our baseline condition, the task elicited increased arousal which modulated M1 activity. Therefore, in this case we cannot claim that general motor resonance is an indicator of action understanding. However, measuring grip-specific changes by employing a two actions/ two muscles design provides insights into the muscle-specific computations taking place in M1 during the processing of different actions.

### Combining kinematic and context information during action observation

In accordance with previous research, grip-specific facilitation patterns for whole-hand versus precision grips were observed in the FDI and ADM muscles when participants observed how these actions unfolded (Alaerts, Senot, et al., 2010; Alaerts et al., 2009; Alaerts, Swinnen, & Wenderoth, 2009b; Koch et al., 2010). We also confirmed across two independent studies that informative cues are sufficient to facilitate the observer’s motor system (de Beukelaar et al., 2016; Kilner et al., 2004; Villiger, Chandrasekharan, & Welsh, 2011). Importantly, we demonstrated that this cue-dependent facilitation is grip specific (de Beukelaar et al., 2016) and persists even if the movement kinematics are occluded from view. Neurophysiological studies in humans and non-human primates support this idea by showing that mirror activity is influenced by non-kinematic triggers (Avanzino et al., 2015; Lagravinese et al., 2017; Lehner, Meesen, & Wenderoth, 2017; Maranesi, Livi, Fogassi, Rizzolatti, & Bonini, 2014; Saucedo Marquez, Ceux, Wenderoth, & Avenanti, 2011) and is still present when well-known actions are occluded from view (Umiltà et al., 2001).

Moreover, our simple Bayesian model made another important prediction, i.e. combining kinematic information with contextual information should evoke slightly stronger motor resonance than context information alone (note that contextual information was correct for only 96% of the trials, thus it was slightly more uncertain than the kinematic information). Evidence in support of this account comes from our combined results from Experiments 1 and 2 showing that grip-specific muscle-facilitation was significantly stronger when both informative cues and full-view movement kinematics are present compared to when only informative cues are accessible. Previous research has suggested that goals and outcomes of an observed action might be represented by a hierarchically organized network of areas in parietal and frontal cortex (Avenanti, Paracampo, Annella, Tidoni, & Aglioti, 2017; Hamilton & Grafton, 2006). Our study revealed physiological evidence that different aspects of action representations do ultimately converge onto M1 as predicted by hierarchical models of predictive coding (see Kilner et al. 2007).

In the predictive coding framework contextual information or prior knowledge is able to guide the observer towards inferring the most likely intention of an observed movement even if sensory input is uncertain or lacking completely. This theory is supported by several studies showing that predictions regarding observed actions can modulate the mirror responses (Cross, Stadler, Parkinson, Schütz-Bosbach, & Prinz, 2013; Plata Bello, Modroño, Marcano, & González-Mora, 2015; Press, Weiskopf, & Kilner, 2012; Thioux & Keysers, 2015; Wurm & Schubotz, 2012, 2017). Our results show that grip specific facilitation of human M1 is consistent with this idea, suggesting that contextual cues activated top-down mechanisms which facilitated highly specific motor representations in M1. This suggests that M1 might be an important area for action understanding, which also represents information other than the observed kinematics. This information is likely to be mediated by parietal and frontal regions which are part of the MNS (Chouinard & Paus, 2010; Majdandzc, Bekkering, Van Schie, & Toni, 2009; Toni, Rushworth, & Passingham, 2001) and project directly or indirectly to M1.

Even though our results fully support key predictions of the Bayesian model, one has to note that our experiment varied the uncertainty of kinematic and contextual information according to a categorical rather than a probabilistic regime. Future studies might therefore test contextual cues which vary more strongly in accuracy or might use kinematic information which varies in reliability and directly model the MEP_mod_ data. Moreover, priors often refer to knowledge representing the statistics of a task or the environment which is formed by experience. By contrast, our cues represent contextual priors (Seriès and Seitz 2013, i.e. the relevant contingencies are simple and can be rapidly learned and manipulated). In conclusion, the new insight revealed by our study is that human M1 activity caused by motor resonance during movement observation reflects kinematic and contextual information as predicted by a classical Bayesian model. M1 activity did not only reflect general arousal effects but could be linked to specific motor representations. Our findings suggest that both high and low-level representations of observed movements converge in motor areas and particularly M1. These representations are likely to be mediated by bottom-up mechanisms that drive motor activity as a function of the observed kinematics, and top-down mechanisms that activate motor representations associated with arbitrary cues. Our findings suggest that M1 is an important neural substrate for mediating action understanding which integrates kinematic and non-kinematic information in accordance to a predictive coding model.

## ACKNOWLEDGEMENTS

This project was supported by an SNSF Grant (320030_175616).

